# Stochastic transcription in the p53-mediated response to DNA damage is modulated by burst frequency

**DOI:** 10.1101/679449

**Authors:** Dhana Friedrich, Laura Friedel, Andreas Herrmann, Stephan Preibisch, Alexander Loewer

## Abstract

Discontinuous transcription has been described for different mammalian cell lines and numerous promoters. However, our knowledge of how the activity of individual promoters is adjusted by dynamic signaling inputs from transcription factor is limited. To address this question, we characterized the activity of selected target genes that are regulated by pulsatile accumulation of the tumor suppressor p53 in response to ionizing radiation. We performed time resolved measurements of gene expression at the single cell level by smFISH and used the resulting data to inform a mathematical model of promoter activity. We found that p53 target promoters are regulated by frequency modulation of stochastic bursting and can be grouped along three archetypes of gene expression. The occurrence of these archetypes cannot solely be explained by nuclear p53 abundance or promoter binding of total p53. Instead, we provide evidence that the time-varying acetylation state of p53’s C-terminal lysine residues is critical for gene-specific regulation of stochastic bursting.

## INTRODUCTION

Cells constantly respond and adapt to extrinsic and intrinsic stimuli to mediate appropriate cell fate decisions. Intracellular signaling pathways connect these incoming signals to cellular responses through changes in abundance, localization, or post-translational modification of signaling molecules. Recent studies employing time-resolved single cell measurements highlighted that stimulus specific temporal activity patterns contribute to regulating gene expression and cellular phenotypes as well ((Nelson et al., 2004); (Tay et al., 2010); (Batchelor et al., 2011); (Hao and O’Shea, 2011); (Purvis et al., 2012)). Associated transcription factors (TFs) often show pulsatile dynamics with time-scales ranging from seconds (NFAT4) and minutes (NF-kB, Msn2, Erk) to hours (p53), ((Yissachar et al., 2013); (Tay et al., 2010); (Hao and O’Shea, 2011); (Shankaran et al., 2009);(Lahav et al., 2004)). However, it still remains unclear how molecular circuits convert information from pulsatile TF dynamics to distinguishable expression profiles and how pulses of transcription factors quantitatively control transcription rates of target genes at individual promoters.

To address these questions we focused on the tumor suppressor p53. Its main function is to protect genetic integrity and inhibit uncontrolled proliferation in the context of cellular stress and transformation. In unstressed cells, p53 nuclear abundance is kept low through ubiquitination by the ubiquitin ligase MDM2 and rapid proteasomal degradation ((Haupt et al., 1997); (Kubbutat et al., 1997)). In response to ionizing radiation (IR) induced DNA double strand breaks (DSBs), p53 accumulates in a series of undamped pulses ((Lahav et al., 2004); (Batchelor et al., 2008)) (Fig 1A). In contrast, other insults such as UV radiation or chemotherapeutic drugs lead to sustained accumulation of the transcription factor ((Batchelor et al., 2011);(Paek et al., 2016)). P53 dynamics contribute to determining cellular outcomes, as pulsatile p53 accumulation is correlated with transient cell fate programs (cell cycle arrest), while sustained p53 levels induce terminal responses (apoptosis, senescence) (Purvis et al., 2012). To enable stimulus dependent regulation of cellular phenotype, p53 activates the concerted transcription of target genes related to apoptosis, cell cycle arrest, DNA repair or senescence. It has been shown that p53 activation leads to the expression of over 300 directly targeted protein-coding genes and noncoding RNAs (Fischer, 2017). However, for many targets the quantitative relation between p53 levels and transcriptional output has not been described. Moreover, p53 has also been detected at target sites in absence of DNA damage despite low nuclear abundance ((Nikulenkov et al., 2012); (Younger and Rinn, 2017)).

**Figure 1.**
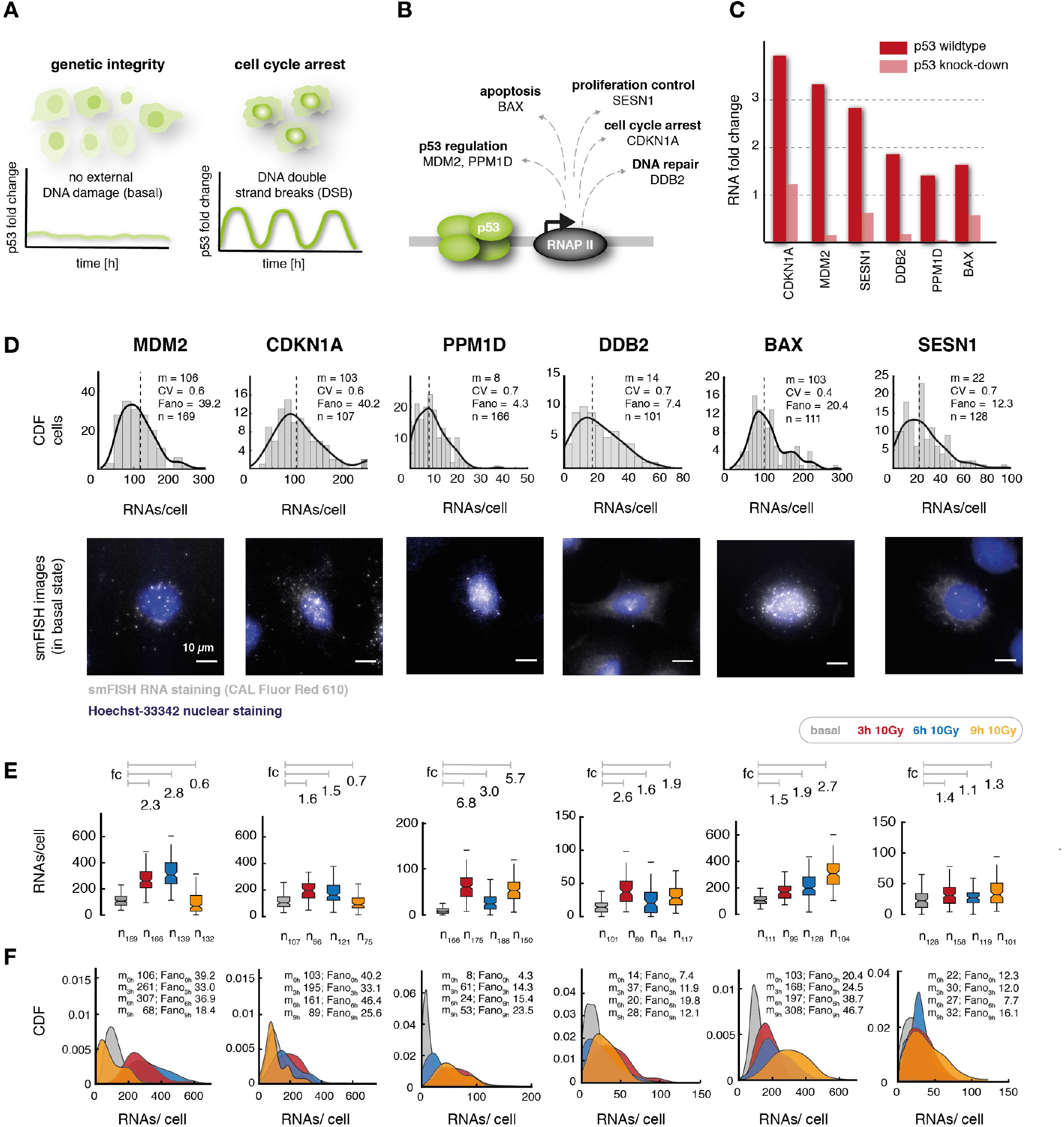
Single cell quantification of p53 dependent transcription highlights distinct patterns of gene expression upon DNA damage. **A** p53 has been show to response with a series of undamped pulse to **γ**-irradiation leading to cell cycle arrest while intrinsic DNA damage during cell cycle does not induce regular pulsatile p53 and subsequent gene expression programs. Schematic representation of p53 dynamics in both cellular conditions. **B** We selected p53 target genes that are involved in different cell fate programs ranging from apoptosis (BAX), DNA repair (DDB2) cell cycle arrest (CDKN1A), proliferation control (SES-N1) and the regulation of the p53 network itself (PPM1D and MDM2). **C** The selected p53 target genes show a similar mean fold change of induction after 10 Gy **γ**-IR in RNA-Seq experiments in MCF10A cells at 4 h, while stable shRNA knock-down of p53 strongly decreases target gene expression, highlighting their dependence on p53 activity for transcriptional up-regulation after DNA damage **D** smFISH staining and quantitative analysis of p53 targets shows a broad variability of RNA counts per cell for all genes in basal conditions. *Upper panel*: Histogram of quantitative analysis of RNAs per cell for each target gene before (basal) DNA damage. Dashed line: median; solid line: fit cumulative distribution function of the dispersion (CDF), m: median, CV: coefficient of variation, Fano: Fano factor; *lower panel*: Fluorescence microscopy images of smFISH probes CAL-Fluor 610 (grey) and Hoechst 33342 (blue) staining overlayed, scale bar corresponds to 10 μm distance, images were contrast and brightness enhanced for better visualization. **E** We quantified RNAs per cell for each target gene before (basal, grey) and 3 h (red), 6 h (blue) and 9 h (orange) after DNA damage (10 Gy IR). smFISH based single cell analysis of gene expression patterns highlights distinct RNA counts for p53 targets. RNA counts per cell are displayed as boxplots. Whisker show 25th, 75th percentile, n: number of analyzed cells, fc: median fold of induction relative to time point basal (indicated by grey lines) **F** Despite a clear change in median levels (m: median), single cell analysis displays a strong dispersion that overlaps for the different conditions, as shown by the strongly overlapping distributions of RNA counts per cells, represented as cumulative distribution function (CDF). This is further highlighted by high Fano factors (Fano).

According to the affinity model, the susceptibility of a target gene promoter to p53 dependent gene expression is defined by the sequence of the corresponding p53 response element (RE). In this model, genes inducing transient phenotypes such as cell cycle arrest tend to have higher affinity for p53 binding compared to genes inducing terminal cell fates such as apoptosis ((Qian et al., 2002); (Weinberg et al., 2005); (Kracikova et al., 2013); (Murray-Zmijewski et al., 2008)). p53 REs consist of two decamers that can be separated by short spacers. Binding site affinity is primarily defined by the central conserved core motif CWWG and the length of the spacer ((Verfaillie et al., 2016); (Riley et al., 2008)). At promoters, p53 has been shown to be involved in a set of key-regulatory mechanisms, including recruitment of histone variants, histone methyltransferases (HMTs), histone acetyltransferases (HATs) and components of the pre-initiation complex (PIC) ((Murray-Zmijewski et al., 2008); (Flores et al., 2002); (Samuels-Lev et al., 2001)). Surprisingly, similar p53 levels can lead to differential locus- and stimulus-specific PIC assembly. Recent live-cell measurements of transcription at the CDKN1A promoter suggested that C-terminal acetylation state instead of p53 abundance is the primary driving factor of transcriptional activation (Loffreda et al., 2017). Even though these mechanisms have been studied in biochemical assays for a selection of p53 targets, our mechanistic understanding of p53’s regulatory role at promoter sites in single cells remains ambiguous at best. Mechanistic studies to date neither include temporal changes in p53 nuclear abundance, nor compare transcriptional activity at individual promoters for more than one target gene. Therefore, our current understanding on how damage induced dynamics of p53 are decoded on the level of gene expression remains limited.

In this study, we aimed to quantitatively measure p53 dependent target gene expression at individual promoters in single cells. We chose a set of well known p53 target genes that represent different cellular response mechanisms as a paradigm and quantified corresponding nascent and matured RNA molecules by single molecule fluorescence in-situ hybridization (smFISH). With the resulting quantitative data, we informed a mathematical model of promoter activity (Bahar Halpern et al., 2015b), which allowed us to extract transcription parameters with single cell and single molecule resolution. Using this approach, we provide a quantitative analysis of stochastic p53-dependent gene expression at defined time points during the DNA damage response to IR induced DSBs and reveal archetypes of p53-mediated expression dynamics. We modulated p53 dynamics using small molecule inhibitors and measured the contribution of its nuclear abundance on promoter activity. Using this approach we found that acetylation in p53’s C-terminal lysine residues is substantially affecting stochastic transcription of target gene promoters.

## Results

### Single molecule mRNA quantification reveals heterogeneous expression of p53 target genes upon DNA damage with distinct abundance patterns

To characterize how p53 pulses in response to DNA damage affect transcriptional activity at individual promoters in single cells over time, we selected a set of well characterized p53 targets involved in different cell fate programs (Fig 1B). The selected genes vary in ***cis***-regulatory architecture, position and sequence of p53 REs (Fig EV1A), but show expression changes in the same order of magnitude 4 h after IR in population studies by mRNA-Seq (Fig 1C). To quantify p53 dependent transcription at individual promoters, we performed smFISH ((Bertrand et al., 1998); (Raj et al., 2008)) in the small cell lung carcinoma cell line A549, which shows characteristic pulses of p53 in response to IR ((Finzel et al., 2016); (Stewart-Ornstein and Lahav, 2017)) (Appendix Fig S1A,B). We assigned mRNAs to their cells of origin, using simultaneous nuclear and cytoplasmic staining and enumerated mRNAs at their subcellular localization using custom analysis scripts in combination with FISH-quant (Appendix Fig S2) ((Carpenter et al., 2006); (Mueller et al., 2013)) (see Methods section). To determine required sample sizes, we analyzed the reproducibility of our quantitative data on biological replicates of MDM2 datasets (Appendix Fig S3).

Surprisingly, our analysis showed that all selected targets were transcribed with considerable RNA counts in absence of DNA damage (Fig 1D). Basal mRNA levels varied from a few molecules to several hundreds. For all target genes, we also observed strong heterogeneity between individual cells (Fig 1D). Recent literature suggested a correlation of cell cycle state and cellular volume with mRNA expression levels in single cells as well as passive buffering of expression heterogeneity through compartmentalization by limiting nuclear export ((Padovan-Merhar et al., 2015); (Battich et al., 2015); (Bahar Halpern et al., 2015a); (Stoeger et al., 2016)). Therefore, we determined the ratio of the Fano factor in the nucleus and the cytoplasm (Fano_nuc_/ Fano_cyt_) for our gene set (Fig EV1B). In agreement with recent work by Hansen et al. ((Hansen et al., 2018)) we see a trend towards noise amplification in cytoplasmic compared to nuclear fractions instead of attenuation (Figs EV1B, Appendix Fig S4). We also only observed a minor contribution of cell cycle and volume to heterogeneity (as measured by coefficient of variation, CV) (Appendix Fig S5).

To analyze how RNA counts, localization and variability evolve during the pulsatile p53 response in individual cells, we measured target gene mRNAs in single cells at selected time points after IR covering p53 dependent activation of transcription, its adaptation and progression after reinitiation by upstream kinases ((Lahav et al., 2004); (Batchelor et al., 2008)). In A549 cells, these time points correspond to basal (undamaged), 3 h post 10 Gy (1st p53 peak), 6 h post 10 Gy (minimum after 1st p53 pulse) and 9 h post 10 Gy (2nd p53 peak) (Appendix Fig S1B). To validate pulsatile p53 level in A549 *wild type* cells, we performed quantitative measurements based on immunofluorescence staining (Appendix Fig S6). Although an increase in the heterogeneity of p53 dynamics from the first to the second pulse was detected, our measurements indicate sufficient synchrony in A549 cells until 9 h after 10 Gy IR. In agreement with previous work, our smFISH based analysis showed that p53 target genes were expressed in different patterns over time with similar mean induction (fc) during the first p53 pulse for most target genes except PPM1D and gene specific changes at later time points (Fig 1E) ((Porter et al., 2016); (Hafner et al., 2017); (Hanson et al., 2019)). We also measured changes in the distribution of mRNA counts for each individual target when DNA damage is applied (Fig 1F) and observed gene specific shifts in the variability of RNA counts (Fig EV1C, Appendix Fig S5), indicating mechanistic changes in p53 dependent transcription upon DNA damage.

### Single cell characterization of promoter states shows frequency modulation of bursty transcription

At an individual promoter, transcription can be either continuous or a stochastic process with episodic periods of active bursting and silent promoter states ((Raj et al., 2006); (Singh et al., 2010); (Zenklusen et al., 2008); (Suter et al., 2011); (Dar et al., 2012); (Golding et al., 2005); (Coulon et al., 2013)). To investigate if p53 target genes encounter bursty or continuous transcription, we quantified the dispersion of mRNAs for all analyzed p53 targets in A549 cells. We observed that the corresponding distributions deviated from Poisson-like dispersions expected for constitutively active promoters (Fano_mRNA_ ≫ 1; Fano_pois_ = 1) (Fig EV1B,C) ((Dar et al., 2016); (Singh et al., 2012)). Single cell mRNA measurements of p53 targets therefore suggest stochastic transcription under basal and induced conditions ((Peccoud and Ycart, 1995); (Kepler and Elston, 2001)). Despite high nuclear RNA counts for some targets such as BAX, MDM2 and CDKN1A, the nuclear to cytoplasmic ratio of mRNAs did not change for most of the analyzed targets upon IR (Appendix Fig S7), indicating that nuclear export is not limiting at this time scale. On the level of promoter activity, RNA numbers per cell can rise by more frequent promoter activation (burst frequency) or a higher rate of transcription during active periods (burst size) (Fig 2A,B) ((Raj et al., 2008); (Larson et al., 2011); (Lionnet and Singer, 2012)). According to the *random telegraph* model an increase in mean mRNA expression via higher burst size leads to an increase in gene expression noise (measured as CV^2^), while an increase through burst frequency correlates with reduced gene expression noise ((Peccoud and Ycart, 1995); (Kepler and Elston, 2001)). We analyzed the CV^2^ versus mean relationship for all p53 targets and observed a trend to attenuated or reduced noise with increasing mean RNA counts 3 h after 10 Gy (Fig EV1D). At later time-points we detected more gene specific correlations (Fig EV1D). In general, an increase of mean mRNA counts upon p53 activation led to reduced or similar gene expression noise compared to undamaged cells (Figs EV1C,D). These results point towards a change in burst frequency rather than burst size ((Dar et al., 2016); (Singh et al., 2012)) and suggest promoter specific regulation at later time points.

**Figure 2.**
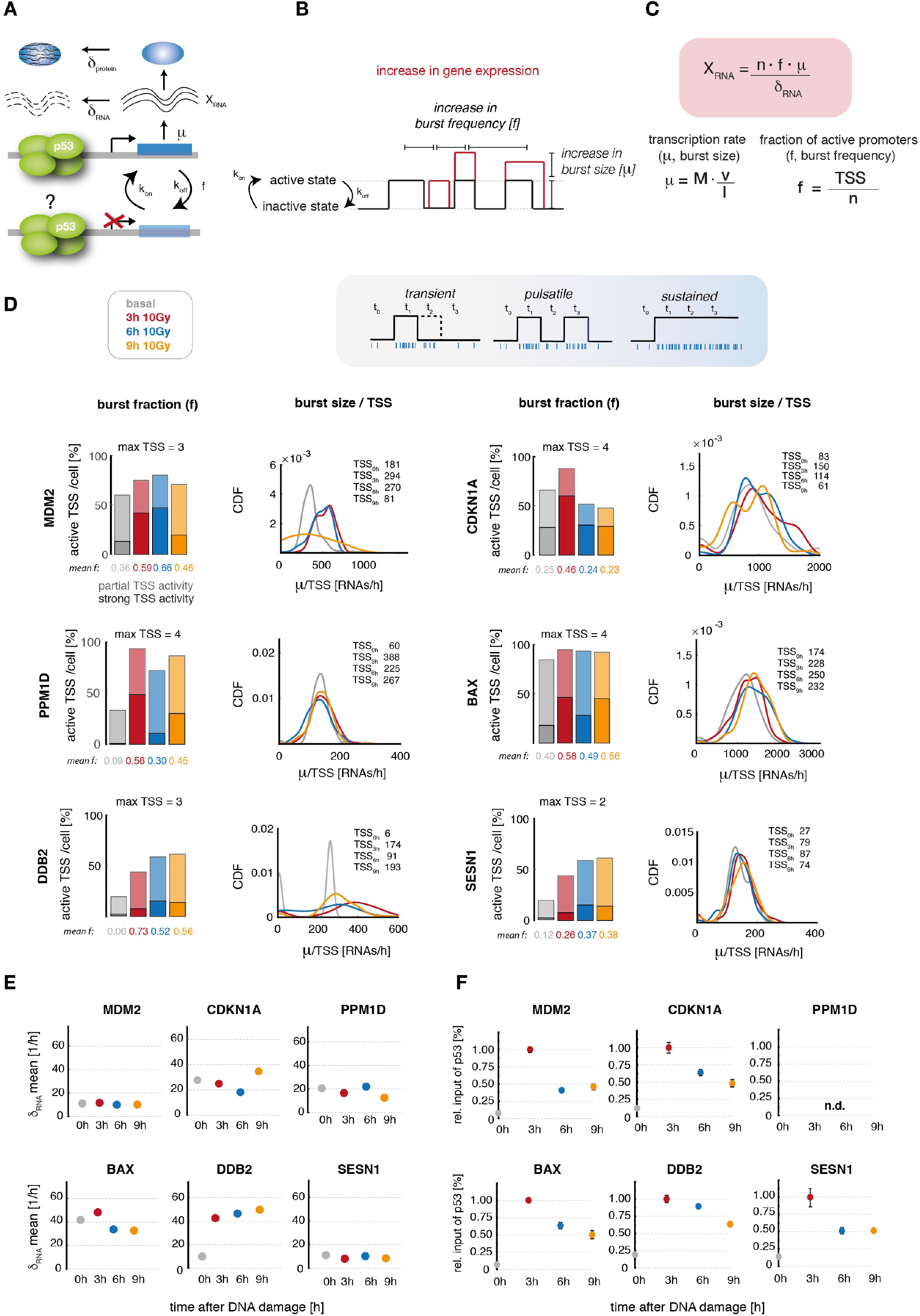
smFISH based analysis at the first and second p53 pulse after IR leads to gene specific stochastic bursting. **A** Schematic illustration of the life-cycle of an mRNA and the rate constants that influence RNA abundance due to stochastic bursting according to previously published models of promoter activity. While <u>burst frequency</u> (f) describes the switching of a promoter between a transcriptionally active and inactive state with the rate constants k_on_ and k_off_, the <u>burst size</u> [μ] describes the rate of RNA transcription in an active period. Additionally, degradation (δ) further influences RNA levels by reducing the cytoplasmic RNA pool. **B** Illustration of promoter activity according to the *random telegraph* model. An increase in RNA levels per cell can be due to a higher burst frequency (more active promoter periods, a higher rate of transcription initiation), or an increase in burst size (a higher rate of RNA transcription in an active period). Additionally, also mixtures of both scenarios are possible. **C** We used smFISH data to calculated promoter activity based on previously published models. Overview of the calculation of stochastic bursting parameters. X_RNA_: number of quantified RNAs/cell, n: number of genomic loci, f: Fraction of active promoters (burst frequency), μ: burst size per cell [RNA/h], δ_RNA_: RNA degradation rate per cell [1/h], M: Polymerase occupancy [RNAs/h], v: RNAP2 speed (estimated as 3 kb/min), l: gene length, TSS: active TSS at the moment of measurement. Further details can be found in Methods section. **D** p53 target genes show an increase in their fraction of active promoters (relating to burst frequency) while the burst size per TSS remains similar upon DNA damage for all time points. *Left panel*: Fraction of active promoters relative to the number of genomic loci [%], shaded colors: partially active cells, solid colors: cells in which a majority of TSS are actively transcribing, average fractions of active promoters are indicated at the bottom of each panel (*mean f*); *Right panel:* relative burst size/TSS: TSS activity divided by the number of active TSS shown as cumulative distribution functions (CDF). Based on promoter activity, we grouped target gene promoters into three archetypes of activity, that are illustrated by graphs (center). **E** Mean degradation rates of target RNAs in transcriptionally active cells as calculated from smFISH data indicate that RNA stability is not changing in the measured time frame upon DNA damage. The plot displays the average RNA degradation rate per cell [1/h] over time after DNA damage, calculated from model (A) in actively transcribing cells for each gene. **F** Relative % input of total p53, indicating binding to target gene promoters after DNA damage. Measured by ChIP and normalized to 3 h after DNA damage for better comparability at later stages in the response. Means and standard deviation from at least three independent experiments are indicated.

As variability in mRNA levels could have other sources than bursty transcription such as differences in stability, we aimed to measure transcription states unambiguously in single cells after IR. Previous work has shown that dual-color labeling of introns and exons by smFISH in combination with mathematical modeling allows to quantify transcription rates, promoter states and mRNA life times in fixed cells (Bahar Halpern et al., 2015b) (Fig 2C). Using the same approach, we designed a second library of smFISH probes for each target gene to identify active sites of transcription based on intron/exon co-staining (Appendix Fig S8A). The fraction of active promoters (burst frequency) can hence be calculated as the ratio of co-stained nuclear dots and the expected number of genomic loci, while the rate of transcription (burst size) is inferred from fluorescence intensity of nascent RNAs at active start sites (Fig 2C) ((Raj et al., 2008); (Bahar Halpern et al., 2015b)). In A549 cells, we detected sites of active transcription only inside nuclei as expected. They varied in number and fluorescence intensity, as introns are spliced and degraded co-transcriptionally (Appendix Fig S8A) ((Levesque and Raj, 2013); (Vargas et al., 2011)). As systematic co-localization analysis showed more than two transcriptional start sites (TSS) for some p53 targets (Appendix Fig S8B), we validated the maximal number of genomic loci for a representative gene independently in A549 cells by DNA-FISH (Appendix Fig S9, Table S1).

To analyze how stochastic bursting at target gene promoters changes with pulsatile p53 after IR, we characterized the fraction of active promoters, RNAP2 occupancy (M), burst size (μ) [RNAs/h] and RNA stability as degradation rate (δ_RNA_) [1/h] (Figs 2D and EV2). For all p53 target genes we detected a strong increase in the fraction of actively transcribing promoters with the first p53 pulse. When p53 levels decreased to basal level at 6 h, we saw that the MDM2, BAX, DDB2 as well as to a lesser extent the SESN1 promoter retained high burst frequencies. In contrast, we detected a lower number of CDKN1A and PPM1D transcription sites. Interestingly, p53 accumulation during the second pulse was not linked to an up-regulation in burst frequency for all targets. RNAP2 occupancy and relative burst size per TSS did not change strongly upon IR for all time-points after 3 h. Furthermore, we did not observe noticeable changes in RNA stability upon IR for the selected genes and time-points upon IR (Fig 2E).

To help our understanding of the observed gene-specific time-dependent patterns of stochastic gene expression, we defined the three promoter archetypes “*sustained”, “pulsatile” and “transient”* and assigned our set of target genes gradually along this spectrum (Fig 2D). For some genes, this resulted in a clear classification, as PPM1D, for example, showed obviously pulsatile promoter activity. For other genes, the assignment was more ambiguous, although they mostly trended towards one archetype of activity.

Transcriptional burst frequency can be modulated by concentration sensitive transcription factor binding ((Senecal et al., 2014); ƒ; (Kafri et al., 2016)), interaction with distal *cis*-regulatory elements (Fukaya et al., 2016), and the H3K27ac state of promoters (Nicolas et al., 2018). To test wether gene specific differences in transcriptional activity can be explained by differential p53 binding or acetylation state, we performed ChIP experiments for selected target genes. H3K27ac remained at high levels at the measured time points without notable differences (Appendix Fig S10). P53 promoter binding reached a maximum at the first accumulation pulse as expected (Fig 2F). Surprisingly, it was not reduced to basal levels at 6 h. Instead, we found that for all analyzed promoters, p53 binding decreased gradually to intermediated levels, although its global concentration varied significantly between the trough and the second peak at 9 h.

### P53 dynamics affect stochastic transcription

Our results so far suggested a gene specific shift in p53’s potency as a transcriptional activator after IR despite continuous promoter binding. As previous work has correlated stimulus dependent p53 dynamics with cell fate specific gene expression (Purvis et al., 2012), we investigated how modulation of p53 dynamics after the first peak affects bursting kinetics of the observed target gene archetypes. To this end, we used small molecular inhibitors in combination with IR to tune the p53 response into transient or sustained dynamics and tested four representative targets from our gene set: MDM2, BAX, PPM1D and CDKN1A.

First, we generated a transient p53 response with only one accumulation pulse using the Chk2-inhibitor BML-277 at 4 h after IR (Fig 3A). This allowed us to focus on gene specific differences during the second p53 pulse, leaving the initial DNA damage regulation of p53 and transcriptional activation of targets unchanged. Our analysis revealed that both PPM1D (resembling *pulsatile* promoter activity) and BAX (resembling *sustained* promoter activity) had reduced burst frequencies when the p53 response was transient, while burst size remained similar (Fig 3B,C; Fig EV3A,B). A direct comparison at the 9 h time point shows that in Chk2 inhibitor treated cells the fraction of active BAX TSS is strongly reduced compared to pulsatile p53, while we see a weaker effect at the PPM1D promoter with a reduction. This indicated that the reoccurrence of a second p53 pulse is necessary to keep those genes in an active transcription mode after the first pulse. Notably, target gene expression is decreased significantly for genes that showed a trend to *transient* promoter activity as well when further p53 pulsing is prevented (Fig EV3A,B). Next we asked how persistent nuclear p53 accumulation affects stochastic bursting. To test this, we used an increasing sequence of the small molecule MDM2 inhibitor Nutlin-3 (Vassilev et al., 2004) after IR to change p53 dynamics from a pulsing to a sustained regime (Fig 3D) (Purvis et al., 2012). Upon Nutlin-3 treatment, burst frequency at the 9 h time point increased for all targets (Fig 3E,F; Fig EV3C,D) including MDM2 and CDKN1A that resembled the *transient* promoter archetype when p53 was pulsing. Interestingly, when p53 was kept at high levels for extended time periods, we did not solely detect an increase in burst frequency, with an increase in the fraction of active promoters of 2.1 fold for CDKN1A and 1.9 fold for MDM2, but also an increase in burst sizes over time, that was >2 fold higher than in response to pulsatile p53 (IR only (Fig 3E,F; Fig EV3C,D). This indicates that sustained nuclear p53 leads to a mechanistic shift in promoter regulation for targets with *transient* promoter activity via a different mechanism than upon IR only treatment. When we compared relative p53 binding under transient and sustained p53 conditions by ChIP, we further detected an increase at all analyzed promoters for sustained p53 (BAX, CDKN1A and MDM2) (Fig EV3E). Notably, transient p53 accumulation upon Chk2 inhibition did not lead to a complete loss of p53 binding at these promoters, but to comparable binding profiles as pulsatile p53 (Fig EV3E).

**Figure 3.**
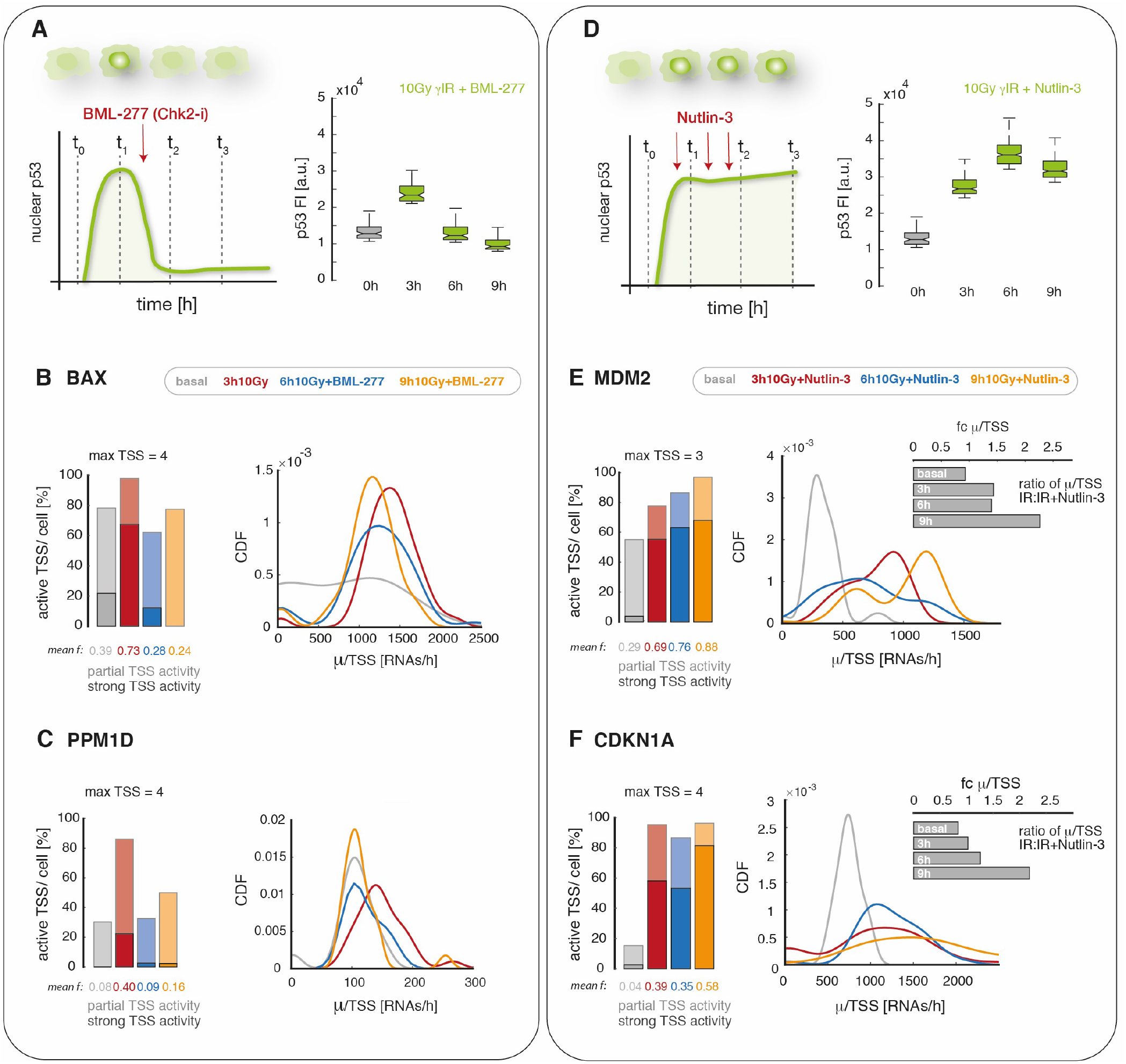
Promoter archetypes change upon modulation of p53 dynamics through small molecule inhibitors. **A** Chk2 inhibition with the small molecule BML-277 induces transient p53 dynamics with only one pulse after 10 Gy IR. A schematic illustration of the experimental setup and quantification of p53 levels in A549 *wild type* cells after irradiation with 10 Gy IR and addition of 10 μM BML-277 by immunofluorescence staining (see Methods section for details) are shown. **B/C** We quantified promoter activity of BAX (B, sustained archetype) and PPM1D (C, pulsatile archetype) after inhibiting the second p53 pulse by Chk2 inhibition. The relative fraction of active promoters (left panel) was reduced at 6 h and 9 h after 10 Gy; the relative burst size per TSS displayed as cumulative distribution function (CDF, right panel) in RNAs/h was not notably affected (right panel). Shaded colors: partially active cells, solid colors: cells in which a majority of TSS are actively transcribing, average fractions of active promoters are indicated at the bottom of each panel (*mean f*); **D** Sequential treatment with Nutlin-3 converts pulsatile p53 dynamics into sustained nuclear levels. A schematic illustration of the experimental setup and quantification of p53 levels in A549 *wild type* cells after irradiation with 10 Gy IR and sequential treatment with 0.75 μM Nutlin-3 at 2.5 h, with 2.25 μM at 3.5 h and 4 μM at 5.5 h post IR based on immunofluorescence staining (see Methods section for details) are shown. **E/F** We quantified promoter activity of MDM2 (E, transient archetype) and CDKN1A (F, transient archetype) upon sequential Nutlin-3 treatment. The relative fraction of active promoters (left panel) strongly increased, changing transient to sustained archetypes. The relative burst size per TSS displayed as CDF in RNAs/h (right panel) increased as well both compared to basal levels and to previous experiments with pulsatile p53 dynamics (inset, fold change calculated for each time point after IR (fc μ/TSS)). Shaded colors: partially active cells, solid colors: cells in which a majority of TSS are actively transcribing, average fractions of active promoters are indicated at the bottom of each panel (*mean f*).

### The K370/382 methylation-acetylation switch contributes to transient promoter activity during the 2nd p53 pulse

The regulatory potential of p53’s highly unstructured C-terminal domain (CTD) has been in the focus of numerous studies aiming to disentangle its functions in modulating gene expression (Sullivan 2018). It has been shown that post-translational modifications of the CTD play a central role in regulating target gene transcription ((Bode and Dong, 2004); (Sims et al., 2004); (Loffreda et al., 2017)). In particular, acetylation of lysine residues K370, K372/73 and K381/82 by p300/ CBP have been associated with a transcriptionally active state (Fig 4A) (Gu et al., 1997). In contrast methylation of K370, K373 and K382 inhibits target gene expression ((Huang et al., 2006); (Shi et al., 2007)). In absence of DNA damage, repressive methylation marks keep p53 transcriptionally inactive. Induction of DSBs induces a rapid change towards CTD acetylation that allow target gene expression ((Loewer et al., 2010);(Berger, 2010)). To test wether C-terminal acetylation contributes to transient MDM2 and CDKN1A expression during the p53 response, we induced pulsatile, transient and sustained p53 accumulation as described above (see Fig 3) and analyzed p53 acetylation at K370 and K382 by Western blot (Fig 4B). We observed that K382ac levels were higher under sustained p53 conditions compared to pulsatile p53 (Fig 4C), suggesting a stabilization of acetylated p53 due to reduced protein turn-over (Li et al., 2002) or reduced lysine methyltransferase (KMT) activity.

**Figure 4.**
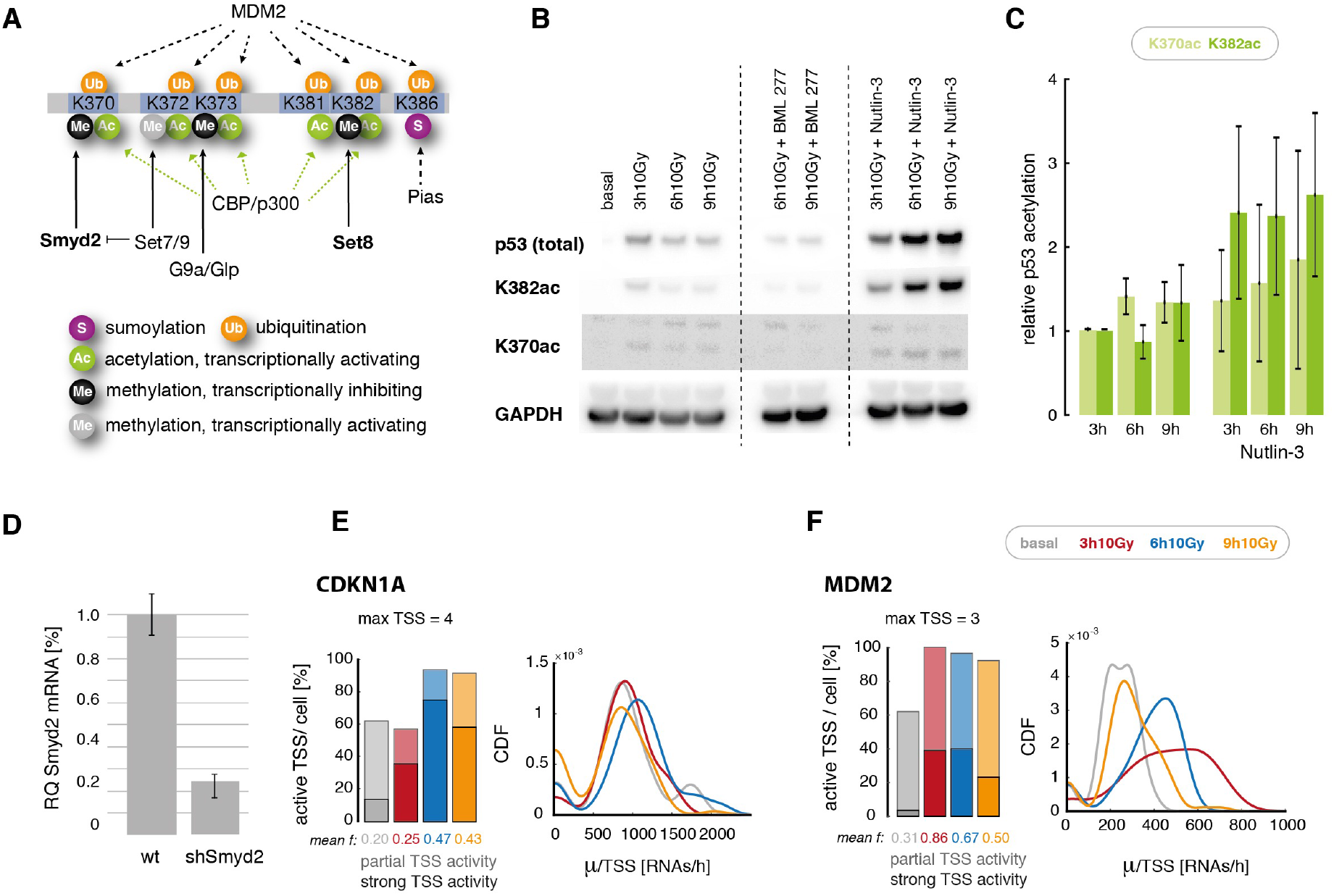
The interplay of p53’s C-terminal lysine acetylation and methylation regulates transiently expressed target genes in response to IR. **A** A schematic illustration of p53’s C-terminal modifications and described functional implications, including key regulatory enzymes. **B** Total p53, p53 acetylated at K382 and K370 as well as GAPDH were measured by Western Blot at indicated time points in the context of different p53 dynamics: pulsing p53 (10 Gy), transient p53 (10 Gy + BML-277, central lanes) and sustained p53 (10 Gy + Nutlin-3, right lanes). See Figure 3 and Methods section for details. **C** The relative change in p53 acetylation at K370 (light green) and K382 (dark green) was quantified from Western Blot and normalized to the total abundance of p53 3h post IR. Means and standard errors from 3 independent experiments are indicated. Acetylation increased over time in the context of sustained p53. **D** The p53-K370 methylase Smyd2 was down-regulated in a clonal stable A549 cell line expressing a corresponding shRNA. Transcript levels were measured in wild type and knockdown cells by qRT-PCR (*see also Figure EV4A*). Mean levels and standard deviation from technical triplicates are indicated. **E/F** Promoter activity of CDKN1A (E) and MDM2 (F) were quantified in Smyd2 knock-down cells. We measured a higher fraction of active promoters [%] after 10 Gray yIR compared to A549 wild type cells (Figure 2), while relative burst size per TSS [RNAs/h] remains unchanged. Shaded colors: partially active cells, solid colors: cells in which a majority of TSS are actively transcribing, average fractions of active promoters are indicated at the bottom of each panel (mean f).

**Figure 5.**
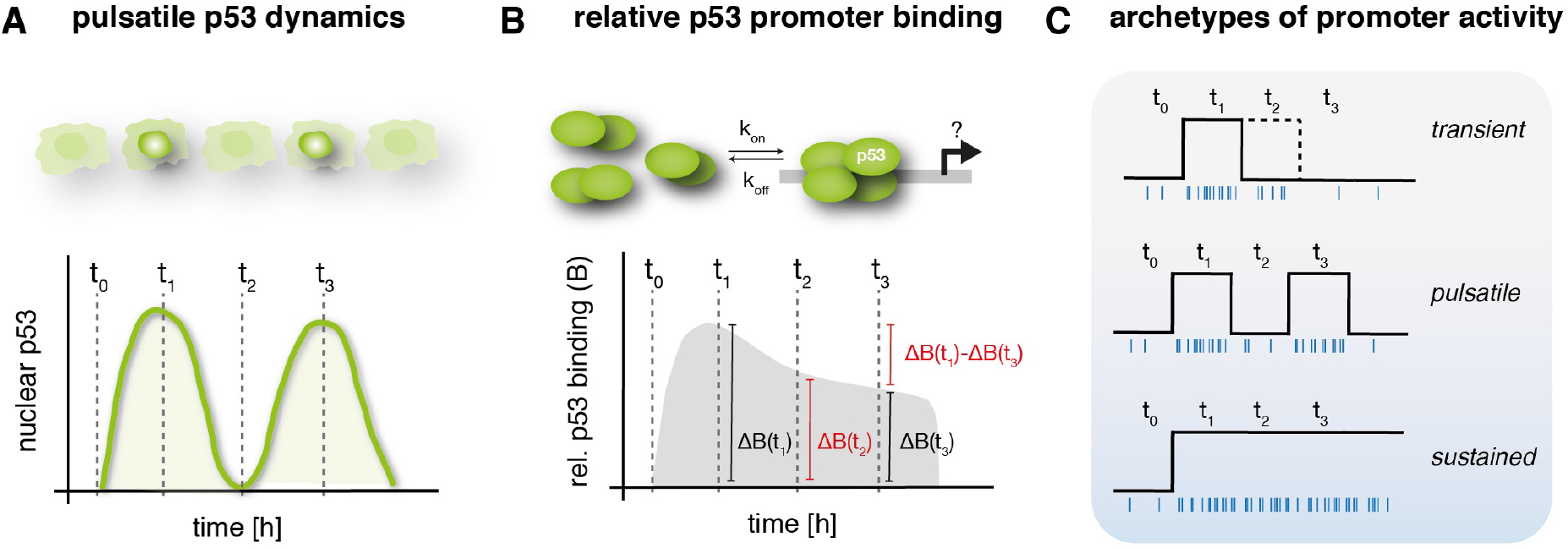
Model of p53 dependent stochastic gene expression. **A** p53 shows reappearing pulses in the nucleus in response to IR that show a high synchrony in A549 cells, as measured by immunofluorescence. **B** p53 promoter binding peaks at 3 h and shows a gradual decrease afterwards, without specific regulation at the time of the second peak. Interestingly, we measured a disconnect between pulsatile p53 in the nucleus and promoter binding at the investigated target genes. **C** p53 target gene expression in response to IR is gene specific with different archetypes of promoter activity that contribute to gene expression profiles inside cells.

Next we asked, how this change in K370/K382 modification state affects stochastic bursting of target genes that we allocated to the *transient* promoter archetype, specifically CDKN1A and MDM2. To this end, we generated stable clonal A549 shRNA knock-down cell lines, reducing the RNA levels of the corresponding methyl transferases Smyd2 and Set8 to 22% and 20%, respectively (Fig 4D). We then characterized burst size and frequency at the same time-points as previously after IR (Fig 4E-F, Fig EV4). While we did not detect strong changes in the fraction of active promoters at basal condition and 3 h after IR compared to A549 *wild type* cells, the mean fraction of active promoters at 9 h was increased from 23% to 43% for CDKN1A and from 46% to 50% for MDM2 in the context of Smyd2 shRNA knock-down (Fig 4E,F) compared to IR irradiated A549 wild type cells. Even though the increase in burst frequency at 9 h after sequential treatment with Nutlin-3 was even stronger and may include also an impact of the change in integrated p53 abundance on bursting, this suggests that Smyd2 mediated methylation contributes to reduced transcription during the second p53 pulse for *transient* p53 targets. Notably, in the context of Set8 knock-down (Fig EV4), we also detected extended expression of *transient* gene CDKN1A through burst frequency modulation, although less prominently than upon Smyd2 knock down. As it has been previously shown that the different lysine residues in p53’s CTD act in concert and embed redundant mechanisms to provide robustness, combinatorial effects of different residues or additional co-factor interaction is likely to lead to transient transcription of MDM2 and CDKN1A.

## DISCUSSION

p53 and other major transcription factors show stimulus specific dynamics correlated with cell fate. While the underlying molecular networks and response mechanisms have been largely characterized, it remains elusive how these proteins regulate gene expression mechanistically at specific promoters in individual cells. In this work, we show that p53 dependent transcription upon IR is intrinsically stochastic and regulated mainly by burst frequency. For our selected panel of p53 targets, we observed that differential regulation of the *on:off* rate of promoter bursting contributes to gene specific dynamics of transcriptional activity. These dynamics could be allocated gradually along a spectrum defined by three archetypes of promoter activity: *transient*, *pulsatile* and *sustained*. These archetypes differed mainly in their response to the second pulse of p53 accumulation upon DNA damage. While target genes resembling the *pulsatile* archetype tended to have low overall expression levels, we could so far not define molecular criteria that would predict expression archetypes for other target genes. Moreover, genes involved in the different response pathways contributed to all archetypes, indicating that the archetype is not directly correlated with cell fate. Further studies of promoter architecture, epigenetic states and combinatorial control of transcription may help to reveal how gene specific modulation of bursting dynamics contributes to structuring the p53 response network upon damage induction.

Frequency modulation of target gene expression has previously been demonstrated for other cellular processes such as c-fos dependent transcription after serum or zinc induction (Senecal et al., 2014), light-controlled transcription by the White Collar Complex (WCC) in Neurospora (Li et al., 2018) and dose-dependent transcriptional regulation by ligand-bound steroid receptors (Larson et al., 2013). Using targeted perturbations it has further been shown that frequency modulation and polymerase pause release are key-regulatory aspects of transcriptional regulation, while RNAP2 recruitment occurs subsequent to burst initiation (Bartman et al., 2019). The simplest model to explain frequency modulation is that the state of a gene is regulated by the *on:off* rate of TF binding to the response element, while the transcription rate in the active state depends on other processes downstream of RE binding. This model suggests that the occupancy of *cis*-regulatory elements by sequence specific TFs can serve as a proxy for transcriptional output (Ptashne and Gann, 2001). Accordingly, we observed coordinated increases in promoter binding and burst frequencies for the initial p53 response to IR and a dependency on recurring p53 accumulation for the *pulsatile* and *sustained* archetypes. However, gene-specific expression patterns at later time points could not be explained by the relatively uniform intermediate binding levels found at all promoters analyzed. Interestingly, we also observed a disconnect between nuclear protein levels and DNA binding after the first pulse of p53 accumulation. Both observations argue against a simplified model of affinity-based regulation of burst frequencies and suggest other regulatory mechanisms during the DNA damage response.

Surprisingly, we observed a gradual decrease in p53 promoter binding after the first accumulation peak instead of a tight coupling to p53 levels even in absence of a second p53 pulse (Fig EV3E). How is p53 stabilized at promoters while total p53 levels are reduced to basal state, despite fast binding kinetics of only a few milliseconds (Loffreda et al., 2017)? As relative binding curves were similar for all target genes, a global increase in DNA binding activity or selective stabilization of chromatin bound p53 can be assumed. For example, it has been previously shown that tetramerization of p53 leads to a stabilization of DNA binding in response to DNA damage (Gaglia and Lahav, 2014). In future studies, it would be interesting to investigate by fluorescence correlation spectroscopy if an increase in the tetrameric p53 population can be observed at 6 h after IR compared to basal state.

Another possibility would be that the promoter associated p53 pool shows dominant PTMs at C-terminal lysine residues that are mutually exclusive with MDM2-dependent ubiquitination. DNA damage induces numerous post-translational modifications of the transcription factor, that lead to a stabilization of p53 levels in the nucleus but fulfill a variety of other functions as well. However, in our ChIP experiments, we only resolved total p53. We show that burst frequency is modulated in response to IR and that p53 network perturbations associated with an increase in K370 and K382 acetylation are correlated with higher burst frequencies and, partially, higher burst sizes at p53 target gene promoters. We can only hypothesize about potential mechanisms that lead to these changes as the function of p53s CTD has been controversially discussed in the literature ((Laptenko et al., 2016); (Sullivan et al., 2018)) and its intrinsically disordered topology allows a variety of functions and interactions (Fuxreiter et al., 2008). The CTD binds DNA in a non-sequence specific manner due to the basic nature of its many lysine residues. This allows sliding along the DNA and promotes and stabilizes the sequence specific binding of the DNA binding domain (DBD) at p53 REs ((McKinney and Prives, 2002); (Laptenko et al., 2015)). Further, it has been shown to interact with many co-regulatory factors that strongly dependent on the post translational modifications state which could additionally affect stochastic bursting. Using perturbation studies we could demonstrate that *transient* expression of CDKN1A and MDM2 are differentially regulated via opposing acetylation and methylation of K370 and K382 residues and can be tuned to different modes of stochastic expression. In line with our findings, a previous study indicated reduced p53 promoter binding and transcription through Smyd2 mono-methylation of K370 (Huang et al., 2006). However, as we still see over 50% p53 promoter binding at 9 h post IR, a reduction in promoter binding mediated through Smyd2 dependent K370me cannot solely explain the *transient* expression of MDM2 and CDKN1A in A549 cells. Moreover, Set7/9 activity leading to inhibition of Smyd2 has been shown to be dynamically regulated during the first p53 pulse after IR (Ivanov et al., 2007). Furthermore, K382 mono-methylation by Set8 induces binding of the chromatin compaction factor L3MBTL1 at CDKN1A and PUMA promoters (West et al., 2010). In contrast, CTD acetylation and DNA binding has been characterized in population studies, leading to controversial results about an increase or decrease in binding affinity ((Gu and Roeder, 1997); (Friedler et al., 2005); (Nakamura et al., 2000)). However, acetylation of C-terminal lysine residues has been linked to its transcriptional activity (Tang et al., 2008). Recently, sophisticated single-molecule studies revealed that transient p53-chromatin interactions are modulated upon activation and interaction times reflect the acetylation state of C-terminal p53 residues (Loffreda et al., 2017). Furthermore, it has also been suggested that an interaction between RNAP2 CTD and disordered regions of transcription factors in nuclear aggregates such as p53’s CTD, can lead to recruitment and transactivation (Sullivan et al., 2018) of RNAP2 into an elongation competent form (Kwon et al., 2013). It is possible to speculate that these mechanisms affect stochastic bursting by a direct or indirect increase in transcription initiation and PIC stability or release of paused RNAP2. However, to our knowledge none of these mechanisms have yet been correlated to repeated pulses of p53 on longer time-scales during the DNA damage response or stochastic bursting at the respective promoters. Notably, it has previously also been suggested that Smyd2 affects the RNAP2 elongation rate independent of p53 (Brown et al., 2006). However, we did not see significant changes in burst size upon Smyd2 knock-down that would be expected from altered RNAP2 elongation rates (Fig 4). Therefore, p53 independent transcriptional inhibition of RNAP2 elongation may only play a minor role in regulation of *transient* p53 targets under our experimental conditions (Brown et al., 2006).

Our data indicate that C-terminal modifications of p53 change between the first and the second p53 pulse. Preventing protein turnover using Nutlin-3 resulted in different promoter regulation and stochastic bursting modalities of p53 target genes, indicating stabilization and accumulation of otherwise transient PTMs. The differences in p53’s first and second pulse activity hint towards a change in upstream processes that re-initiate the p53 response after the first trough. To date, the common view on repeated pulses of nuclear p53 is that ATM and other kinases upstream of p53 are re-activated as long as DNA damage is still present (Batchelor et al., 2008). A change in p53’s PTM patterns may thereby hint towards either another layer of regulation downstream of PI3K-like kinases or other co-regulatory factors that reduce p53 PTMs. Besides C-terminal acetylation, p53 S20 or S46 phosphorylation may also contribute to different archetypes, as both of these modifications correlate with promoter specific binding of p53 after etoposide or actinomycine D treatment of U-2 OS cells (Smeenk et al., 2011).

While we focused on the role of p53 modifications in regulating stochastic target gene expression, other mechanisms have been suggested to control gene-specific promoter activity. For example, long-range enhancer-promoter interactions or forced chromatin looping influence burst frequency in other systems ((Fukaya et al., 2016); (Bartman et al., 2016)) and it has been hypothesizes that enhancer-promoter contacts are necessary for every burst (Chen et al., 2018). A recent study could further show that enhancer-promoter interactions of the Hbb1-1 gene increases burst frequency (Bartman et al., 2019). Histone methylation preserves burst frequency between mother and daughter cells (Muramoto et al., 2012) and histone acetylation can affect transcriptional bursting, mainly burst frequency ((Nicolas et al., 2018); (Suter et al., 2011); (Harper et al., 2011)). Furthermore, nucleosome remodeling has been suggested to be rate limiting for transcriptional activation ((Boeger et al., 2008); (Kim and O’Shea, 2008)). Markers of repressive chromatin architecture, such as CTCF boundaries, cohesine and inhibitory histone marks correlate with inducible expression of p53 targets and have been suggested to play a role in gene-specific dampening of p53 dependent expression upon damage (Su et al., 2015). It will be interesting to investigate in future studies to which extent these mechanisms contribute to regulating gene-specific stochastic transcription of p53 target genes in the response to DNA damage. Interestingly, previous studies have suggested that expression patterns of p53 targets are mainly determined by RNA and protein stability ((Porter et al., 2016); (Hafner et al., 2017); (Hanson et al., 2019)). Based on our model of single cell TSS activity, we see that direct transcriptional regulation of stochastic bursting provides an important contribution as well.

In general our data highlight that p53 pulses allow for a broader diversity in gene specific stochastic transcriptional regulation compared to sustained p53 dynamics, which induces transcription of most p53 targets at high rates. Pulsatile TF nuclear dynamics thereby allow for differential promoter archetypes and fine-tuning of transcription as well as other co-regulatory mechanisms. Besides the pre-dominant hypothesis of robustness of cellular signaling, this may play an important role for expanding the regulatory potential of TFs at target promoters over time.

## MATERIALS AND METHODS

### Cell line and constructs

A549 cells were cultured in McCoy’s medium supplemented with 10% fetal bovine serum, 1% penicillin and streptomycin. When required, the medium was supplemented with selective antibiotics to maintain transgene expression (400 μg/mL G418, 50 μg/mL hygromycine or 0.5 μg/ mL puromycin). We generated lentiviral reporter constructs for p53 using the MultiSite Gateway recombination system (Thermo Fisher Scientific) by fusing the protein coding sequence to the yellow fluorescent protein Venus (YFP) under the control of a constitutive human EF1a promoter. We infected A549 cells with corresponding lentiviral particles together with viruses expressing histone 2B fused to cyan fluorescent protein (H2B-CFP) under the control of UbCp as a nuclear marker. Subsequently, stable clonal cell lines were established and validated. For knock-down p53, we used previously published shRNA vectors to generate Smyd2 and Set8 knock down cells. Therefore we used shRNAs targeting Smyd2 and Set8 using expression of specific oligonucleotides from pRetroSuper.puro as previously described ((Brummelkamp et al., 2002a); (Loewer et al., 2010)). VSV-G pseudotyped retroviral particles expressing SET8 shRNA or p53 shRNA (Brummelkamp et al., 2002b) were produced in 293T cells and subsequently used to infect A549 *wild type* cells. These cells were used as polyclonal populations in further experiments.

### Antibodies and reagents

Stellaris probe-sets for smFISH (Biosearch Technologies) were custom designed for intron and exon regions (see Appendix for oligo list) and conjugated with CAL Fluor 610 (Exons) and Quasar 670 (Introns). We used antibodies against total p53 (FL-393, #6243 and DO-1, #sc-126) from Santa Cruz and against acetylated p53 (K373/382, #ab131442, ab62376) from abcam. A fluorescent labelled secondary antibody conjugated with Alexa Fluor 647 as wells as Alexa Fluor 488 N-Hydroxysuccinimid (NHS, 1-Hydroxy-2,5-pyrrolidindion) and Hoechst-33342 staining solution was purchased from Cell signaling/Life Technologies (Thermo Fischer Scientific, #A-21245,20000). DRB (5,6-dichlorobenzimidazole 1-b-D-ribofuranoside) was purchased from Cayman (used at 10 μM, #1001030250), Chk-2 inhibitor II BML-277 (used at 10 μM) from and Nutlin-3 (used at 0.75 - 4 μM, #N6287) from Sigma.

### Single molecule FISH

A549 cells were cultured for 24 h on 18 mm uncoated coverglass (thickness #1). After treatment cells were washed on ice, fixed with 2% para-formaldehyde (EM-grade) for 10 min at room temperature and permeabilized over night with 70% Ethanol at 4°C. Custom probe sets for sm FISH (Biosearch Technologies) were hybridized at a final concentration of 0.1 μM probe following manufacturers instructions over night at 37°C. Following hybridization procedure, cells were washed and incubated with Alexa Fluor 488 N-Hydroxysuccimid (NHS-AF88) for 10 min at RT for unspecific cytoplasmic protein staining, followed by Hoechst nuclear counterstain. Coverglasses were mounted on Prolong Gold Antifade (Molecular probes, Life technologies). Cells were imaged on a Nikon Ti inverted fluorescence microscope with an EMCCD camera (ANDOR, DU iXON Ultra 888), Lumen 200 Fluorescence Illumination Systems (Prior Scientific) and a 60× plan apo objective (NA 1.4) or using appropriate filter sets (Hoechst: 387/11 nm excitation (EX), 409 nm dichroic beam splitter (BS), 447/60 nm emission (EM); Alexa Fluor 488: 470/40 nm (EX), 495 nm (BS), 525/50 nm (EM); CAL Fluor 610: 580/25 nm (EM), 600 nm (BS), 625 nm (EX); Quasar 670: 640/30 nm (EX), 660 nm (BS), 690/50 nm (EM)). Images were acquired as multi-point of 21 z-stacks of each cell (field of view) with 300 nm step-width using Nikon Elements software. Quantification of RNA counts per cell was performed using FISH-Quant (Mueller et al., 2013) and custom written Matlab software.

### Analysis of smFISH data and quantification of bursting parameters

Multicolor z-stacks from Nikon Element software were extracted into individual tif-stacks and imported into FISH Quant (Mueller et al., 2013). For nuclei and cytoplasmic segmentation two approaches, dependent on the quality of cytoplasmic staining by NHS-AF488 were used. For high-quality cytoplasmic staining and low cell density, the FISH Quant build-in cell profiler interface for automatic cell outline detection was used. Parameters of filtering and local focus projection were optimized per dataset. For dense cells and lower intensity cytoplasmic staining, nuclei were automatically detected in FISH-Quant outline-detection GUI, and cytoplasmic outlines were drawn manually. In both cases each cell outline and nucleus was manually checked for correct segmentation before analysis. TSS were identified based on co-localization of exon and intron signal in nuclei. After identification based on co-localization, we defined the area of a TSS based on the exon signal in all z-planes. In brief, according to the FISH Quant work-flow for spot detection, images were filtered, pre-detection was performed, then spots were fitted and fits were further thresholded to exclude outliers. For TSS detection an average cytoplasmic spot was computed. All analysis was performed using FISH Quant batch processing toolbox. RNA spots counts and respective localization were directly taken from FISH_quant based analysis. Bursting activity was characterized based on previously published models ((Raj et al., 2008), (Bahar Halpern et al., 2015b)). To calculate TSS intensity, we used the FISH Quant parameter TS_Pix_sum (sum of all pixels around brightest pixel of TSS) and the mean intensity of all quantified spots at the respective quantified time-point. Further we calculated correction factor eta for probe position using Trans Quant software (Bahar Halpern and Itzkovitz, 2016), for each gene and corresponding probe set (Appendix Table 2). As correction factor kappa for inferred RNAP2 occupancy we use 1.5 as previously suggested (Bahar Halpern et al., 2015b). As estimated RNAP2 speed we used 50 nt/sec as a range of 6.3 - 71.6 nt/sec has been previously measured in mammalian cell lines (Darzacq et al., 2007). Stacked bar graphs of burst frequency were generated by binning cellular TSS activity into partial and strongly active cells. Bins differ by target gene as follows. SESN1: 1 TSS (shaded), 2 TSS (solid); MDM2: 1-2 TSS (shaded), 3 TSS (solid); CDKN1A: 1 TSS (shaded), >1 (solid), BAX/PPM1D/DDB2: 1-2 TSS (shaded), 3-4 TSS (solid).

### Immunofluorescence

Cells were grown on high precision coverslips #1 and fixed with 2% para-formaldehyde, at the indicated time point after DNA damage. Subsequently cells were permeabilized with 0.1% Triton X-100 (Carl Roth) in phosphate-buffered saline and blocked with 10% goat serum (PAN-Bio-tech). Cells were then incubated with p53-Fl393 for 1 h at 37°C. Cells were washed, incubated with secondary antibody coupled to Alexa Fluor 647 (Cell Signaling), and washed again. Finally, they were stained with Hoechst and embedded in Prolong Gold Antifade (Thermo Fisher Scientific). Microscopy set-up was identical to the above mentioned description for smFISH if not descriptor differently as follows: Images were acquired with a 20× Plan Apo objective (NA 0.75) using appropriate filter sets (Hoechst: 387/11 nm excitation (EX), 409 nm dichroic beam splitter (BS), 447/60 nm emission (EM); Alexa Fluor 647: 640/30 nm (EX), 660 nm (BS), 690/50 nm (EM)). Images were acquired as multi-point datasets. Automated segmentation of nuclei and quantitive analysis of p53 levels based in integrated intensity of the fluorescence signal in each nucleus was performed in Matlab (MathWorks) using custom written software.

### RNA-Sequencing

For RNA sequencing, RNA quality was analyzed with the Agilent RNA 6000 Nano Kit, and the concentration was measured with the Qubit RNA Assay Kit (Invitrogen). Library preparation was carried out with the TruSeq RNA Sample Preparation Kit (Illumina) using barcoded primers. Libraries were sequenced on an Illumina HiSeq using the single read protocol (1 × 100 nt). For analyzing the data, we assembled a list of 400 validated p53 target genes from previously published ChIP- and RNA-seq data ((Nikulenkov et al., 2012);(Menendez et al., 2013)).

### Quantitative Real-Time PCR (qRT-PCR)

mRNA was extracted at the indicated time points using the High Pure RNA Isolation kit (Roche, Mannheim, Germany). cDNA was generated using M-MuLV reverse transcriptase (NEB, Ipswich, MA) and oligo-dT primers. Quantitative PCR was performed in triplicates using SYBR Green reagent (Applied Biosciences) on a CFX96 PCR machine (Biorad). Used Primers were: β-ACTIN forward, GGC ACC CAG CAC AAT GAA GAT CAA; β-ACTIN reverse, TAG AAG CAT TTG CGG TGG ACG ATG; SET8 forward CCC TTC CAC GGG CTG CTA C; SET8 reverse GTG CAG TTT GGT TTG GCA GTT CC; SMYD2 forward CCT CAA CGT GGC CTC CAT GTG; SMYD2 reverse TGG ATG ATC TTT GCC GTG AGC TAC

### DNA FISH

DNA FISH probes were amplified from genomic DNA using custom designed primers. Probes were labelled using DIG-DNA labelling KIT (Roche). For detection five probes were pooled after labelling. Before use, probes were denaturated for 10 min at 70°C and the kept on ice until incubation. Cells were grown on high precision coverslips #1 and fixed with 2% para-formaldehyde, then washed with PBS and 2x SSC following RNAse A incubation for 2 h. Afterwards a 70% formamide shock/2xSSC for 5 min was applied to reduce secondary structures and DNA was denaturated for 10 min at 80°C. Afterwards cells were rinsed in 50% formamide/2xSSC and washed with PBS and incubated with denaturated probe for 72 h in humidified chamber sealed with rubber cement on a hybridization slide (Thermo Fisher). Afterwards cells were washed with 50% formamide/2xSSC at 42°C, 0,1%SCC at 60°C and 4x SSC/0,1% Tween at 42°C and PBS. To detect DIG labelled DNA probes, anti-DIG antibody was used, cell were then stained with Hoechst and mounted in Prolong Gold Antifade (Fisher Scientific). Cells were imaged on a Nikon Ti inverted fluorescence microscope with a ORCA R2 CCD camera (Hamamatsu), Lumen 200 Fluorescence Illumination Systems (Prior Scientific) and a 100× plan apo objective (NA 1.45) using appropriate filter sets (Hoechst: 387/11 nm excitation (EX), 409 nm dichroic beam splitter (BS), 447/60 nm emission (EM); Alexa Fluor 647: 640/30 nm (EX), 660 nm (BS), 690/50 nm (EM)). Images were acquired as single-points of 21 z-stacks of each cell (field of view) with 300 nm step-width using Nikon Elements software. In represented images (SI Figure 3) DNA-FISH stained images were median filtered, self-subtracted, maximum-projected and overlayed with nuclear (Hoechst) staining for visualization purposes using FiJi (Schindelin et al., 2012).

### Chromatin Immuno Precipitation (ChIP)

1.6×10^7^ cells per condition were washed once with PBS and crosslinked with 1% formaldehyde in PBS for 10 min. Cells were rinsed with cold PBS and the fixation was stopped using 125 mM Glycine in PBS for 5 min. Cells were washed with cold PBS and harvested in PBS supplemented with 1 mM PMSF. The cell pellet was resuspended in Lysis buffer (5 mM Tris-HCl, pH 8.0, 85 mM KCl, 0.5% Igepal-CA630 supplemented with protease inhibitor cocktail from Roth and 1 mM PMSF) and incubated on ice for 20 min. The nuclear pellet was collected by centrifugation, resuspended in Sonication buffer (50 mM Tris-HCl, pH 8.1, 0.3% SDS (w/v), 10 mM EDTA supplemented with 1 mM PMSF and Protease Inhibitor Cocktail) and incubated for 30 min on ice. Chromatin was sonicated using the Covaris S220 Sonicator (PIP 105, Duty Factor 2%, CPB 200, 2 min). The sonicated samples were centrifuged and the supernatant collected. 80 μg of chromatin were diluted with Dilution buffer (16.7 mM Tris-HCl, 167 mM NaCl, 0.01 % SDS (w/v), 1.2 mM EDTA, 1.1% Triton (v/v), 1 mM PMSF, Protease Inhibitor Cocktail) and incubated overnight at 4°C with 5 μg p53 antibody (FL-393, Santa Cruz) or a control IgG (Normal rabbit IgG, EMD Millipore). To collect the Immunocomplexes, 25 μl of Dynabeads Protein G (Thermo Fisher Scientific) were added for 2 h at 4 °C. The beads were washed once with Low Salt Washing Buffer (0.1% SDS (w/v), 2 mM EDTA, 20 mM Tris–HCl pH 8.1, 1% Triton X-100 (v/v) and 150 mM NaCl), High Salt Washing Buffer (0.1% SDS (w/v), 2 mM EDTA, 20 mM Tris–HCl pH 8.1, 1% Triton X-100 (v/v) and 500 mM NaCl) and LiCl Washing Buffer (10 mM Tris-HCl pH 8.1, 1 mM EDTA, 1 % IGEPAL CA630 (v/v), 1% Deoxycholic acid (w/v), 250 mM LiCl) and TE-Buffer (10 mM Tris-HCl pH 8.1, 1 mM EDTA). The DNA was eluted from the beads for 30 min at 37 °C in Elution Buffer (1% SDS, 0.1 M NaHCO3) twice. Crosslinks were reversed by adding 200 mM NaCl and by subsequent incubation at 65°C overnight. 50 μg/mL RNase A was added for 30 min at 37°C, then 100 μg/mL Proteinase K, 10 mM EDTA and 40 mM Tris-HCl ph 6.5 were added and the samples were incubated for 3h at 45°C. The DNA was cleaned up using the Monarch® PCR & DNA Cleanup Kit. For qPCR, 3 μl of each sample and the following primers were used: BAX forward: AAC CAG GGG ATC TCG GAA G, BAX reverse: AGT GCC AGA GGC AGG AAG T; MDM2 forward: GTT CAG TGG GCA GGT TGA CT, MDM2 reverse: CGG AAC GTG TCT GAA CTT GA; CDKN1A forward: AGC CTT CCT CAC ATC CTC CT, CDKN1A reverse: GGA ATG GTG AAA GGT GGA AA; DDB2 forward: CTC CAA GCT GGT TTG AAC, DDB2 reverse: CAC AGG TAG CCG AGC TAA G; SESN1 forward: GCC GCG GTC ATG TAA ATG AAA G, SESN1 reverse: GAC TTG TCC AGA CGA CAA TG; RRM2B forward: GCT TGC TGG GAA ATC TTG AC, RRM2B reverse: CTG GTC ACC CAG TTG GAA G; PPM1D forward: CGG ACA AGT CCA GAC ATC, PPM1D reverse: TTC GAC GAC GCC GAG AAG

### Western Blot and Immunodetection

Cells were plated 2 days before experiments in 6 cm dishes at 5×10^5^ cell density. After IR, we harvested cells at indicated time points and isolated proteins by lysis in the presence of protease and phosphatase inhibitors (Roth and Sigma-Aldrich), Trichostatin A (APExBio) and Deacetylase Inhibitor Cocktail (MedChemExpress). Total protein concentrations were measured by Bradford assay (Roth). Equal amounts of protein were separated by electrophoreses on NuPAGE 4-12% Bis-Tris Gels (Invitrogen) and transferred to PVDF membranes (GE Healthcare) by electroblotting (Bio‐Rad). We blocked membranes with 5% bovine serum albumin, incubated them overnight with primary antibody. The next day, membranes were washed, incubated with secondary antibody coupled to peroxidase, washed again, and protein levels were determined using chemoluminescence (Western Bright Quantum, Advansta). Precision Plus Protein Dual Color Standards (BioRad) was used for molecular mass comparison. GAPDH and acetylated p53 were detected on the same membrane. The antibodies were stripped to detect total p53 levels. Blots were quantified using FiJi (Schindelin et al., 2012). Used antibodies were: anti-GAPDH (Sigma Aldrich, G9545), anti-p53 (santa cruz #DO1, Cell Signaling, #9282), anti-p53K70ac (abcam, ab183544), anti-p53K382ac (abcam, ab75754), goat-anti-rabbit-HRP (Thermo Scientific), goat-anti-mouse-HRP (Thermo Scientific).

## ACKNOWLEDGEMENTS

We thank Andrea Grybowski (Max Delbrück Centrum Berlin) and Petra Snyder (Technische Universität Darmstadt) for excellent technical assistance. Furthermore we thank Florian Mueller for great help on the usage of FISH Quant and gratefully acknowledge support of the Light Microscopy Facility of the BIMSB at the Max Delbrück Centrum Berlin. We thank members of the Loewer lab at the MDC Berlin and the TU Darmstadt as well as the Preibisch lab at Max Delbrück Centrum Berlin for helpful discussions during the study and preparing the manuscript. This work was supported by fellowships from the Joachim Herz Foundation (to D.F.) and German Research Foundation (GRK1657 to L.F.) as well as a grant from Einstein Foundation Berlin (to A.H. and A.L.).

## AUTHOR CONTRIBUTIONS

A.L. and D.F. designed experiments and conceived the study. D.F. performed experiments, data analysis and prepared figures. L.F. performed ChIP and Western Blot experiments. S.P. and A.H. provided substantial resources and helped with interpreting data. D.F. and A.L wrote the manuscript with contributions from all authors. A.L. supervised the research.

## TABLES AND THEIR LEGENDS

**Table 1 - Overview of cells, TSS and quantified parameters**

Overview table of quantified cells and transcription sites including the average quantified values of the different parameters.

**Table 2 - smFISH probe sets**

smFISH probes sets, exon probes were labelled with CAL Fluor 610, Intron probes were labelled with Quasar-670.

## EXPANDED VIEW FIGURES

**Figure EV1.**
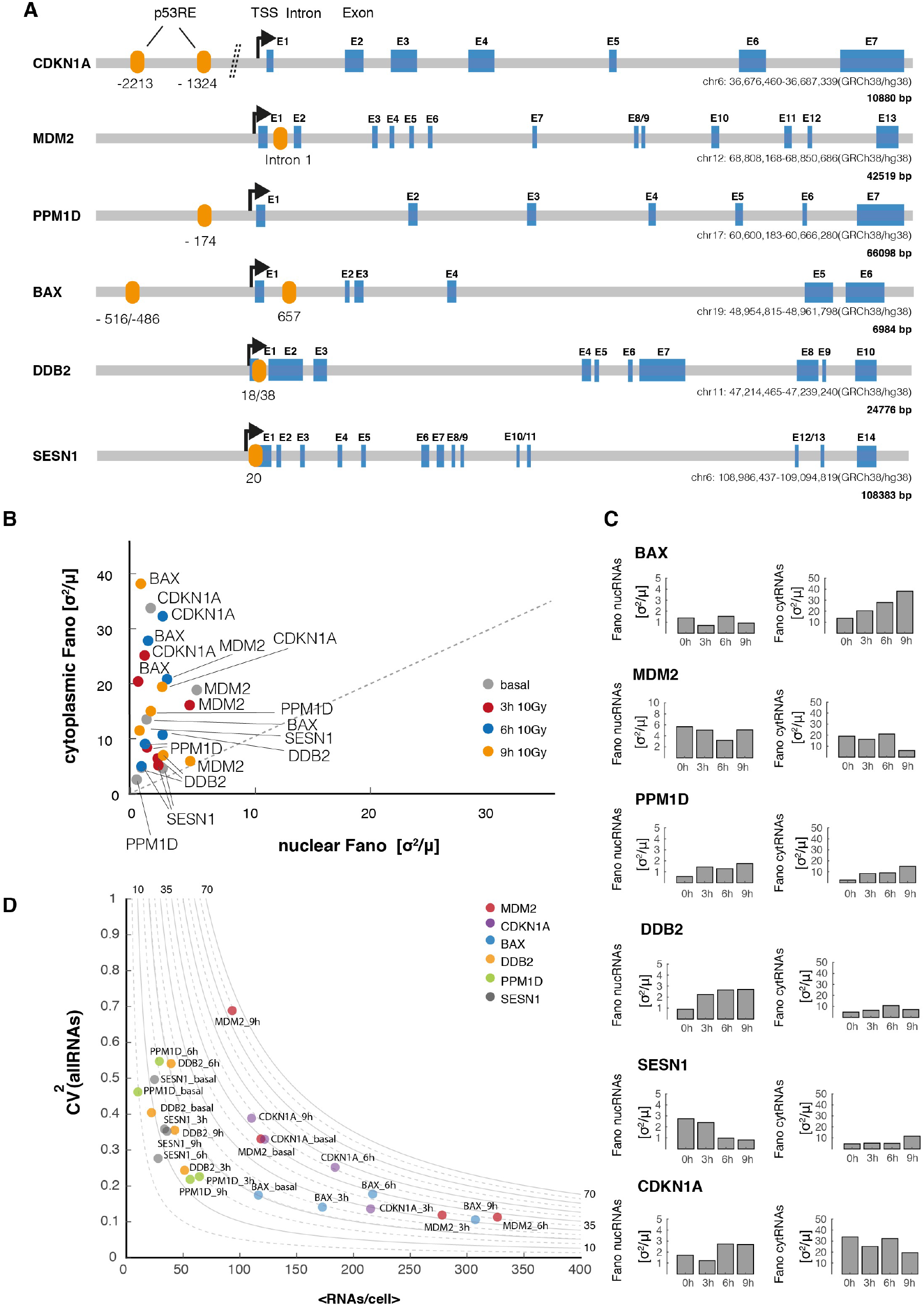
Noise in RNA abundance indicates stochastic bursting and a change in burst frequency after DNA damage. **A** p53 target genes have different genomic architecture and *cis*-regulatory logic. The number and position of p53 response element varies dependent on the target gene. A schematic illustration shows the relative positioning of p53 response elements (RE), transcriptional start sites (TSS) and exons. Gene length and chromosomal positions are indicated. **B** We calculated the Fano factor of RNA counts in the nuclear and cytoplasmic fraction of cells for all p53 targets as a measure of gene expression noise over time after DNA damage 10Gy IR). Data points are labeled, time points are visualized by the indicated color code. The diagonal is shown as a guide to the eye (dashed line). **C** After DNA damage, changes in gene expression noise as measured by Fano factors are gene specific. Fano factors for nuclear (left) and cytoplasmic (right) RNAs are shown for the indicated target genes and time points. Cytoplasmic fractions in general show higher noise levels, but both sub-cellular regions seem to undergo specific changes in noise that do not necessarily follow the same trends. **D** The squared coefficient of variation (CV^2^) in relation to mean RNA count per cell is shown for all p53 target over time after DNA damage (10Gy IR). Data points are labeled with target gene and time points after IR. Hyperbolic lines corresponding to estimated burst sizes based on previously suggested models are shown as guides to the eye ((Dar et al., 2016),(Singh et al., 2010)).

**Figure EV2.**
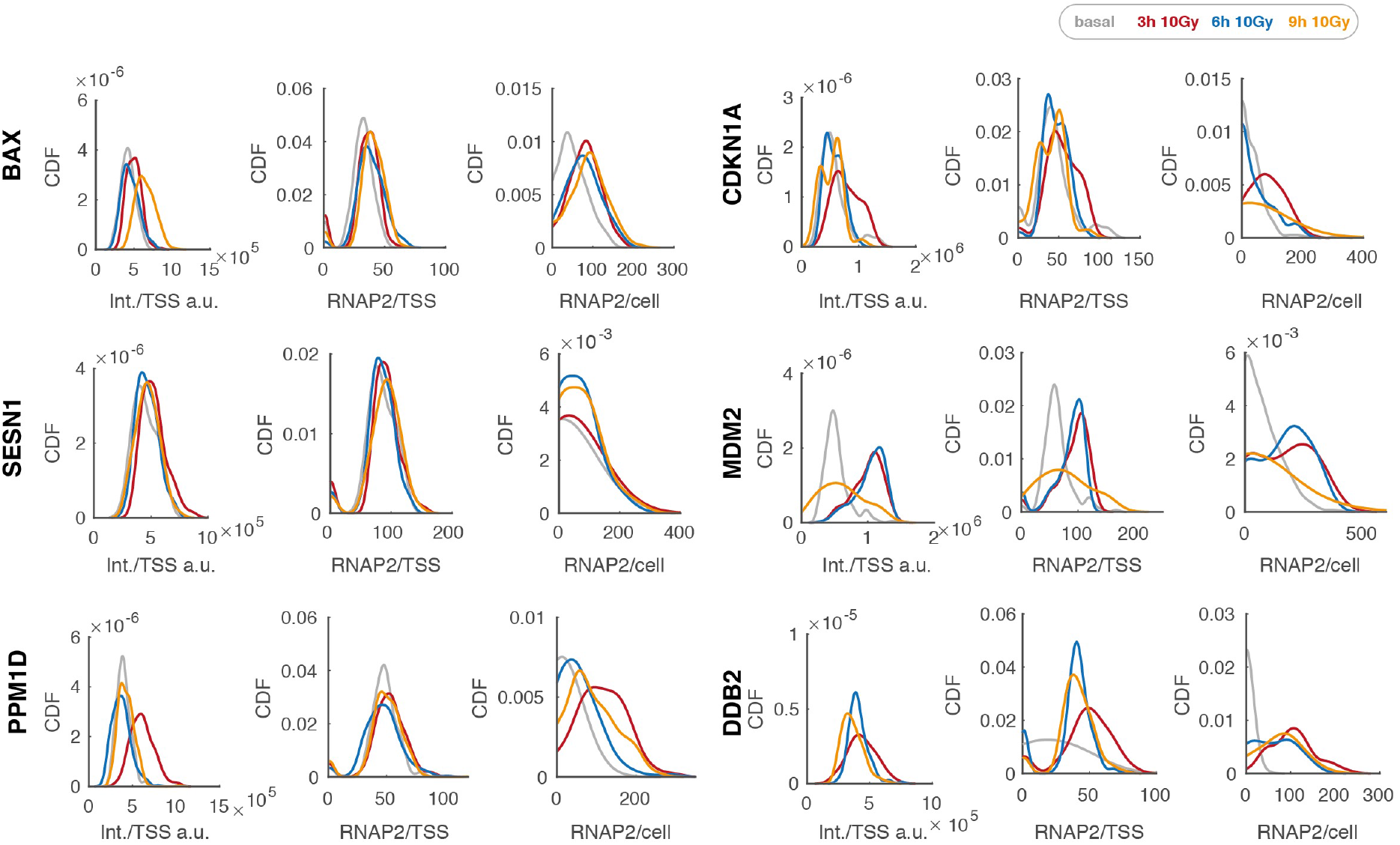
Calculation of RNAP2 occupancy and burst size from TSS intensity. The left panel for each target gene shows the quantified TSS intensities from FISH-Quant displayed as cumulated distribution function of all TSS per cell over time. Center panels indicate RNAP2 occupancies at an individual TSS, right panels the RNAP2 occupancies in the whole cell as calculated from the relative intensity of a TSS and the average cytoplasmic mRNA intensity (see Methods section for details). These occupancies were used to calculate burst sizes per hour. Different time points are visualized by the indicated color code.

**Figure EV3.**
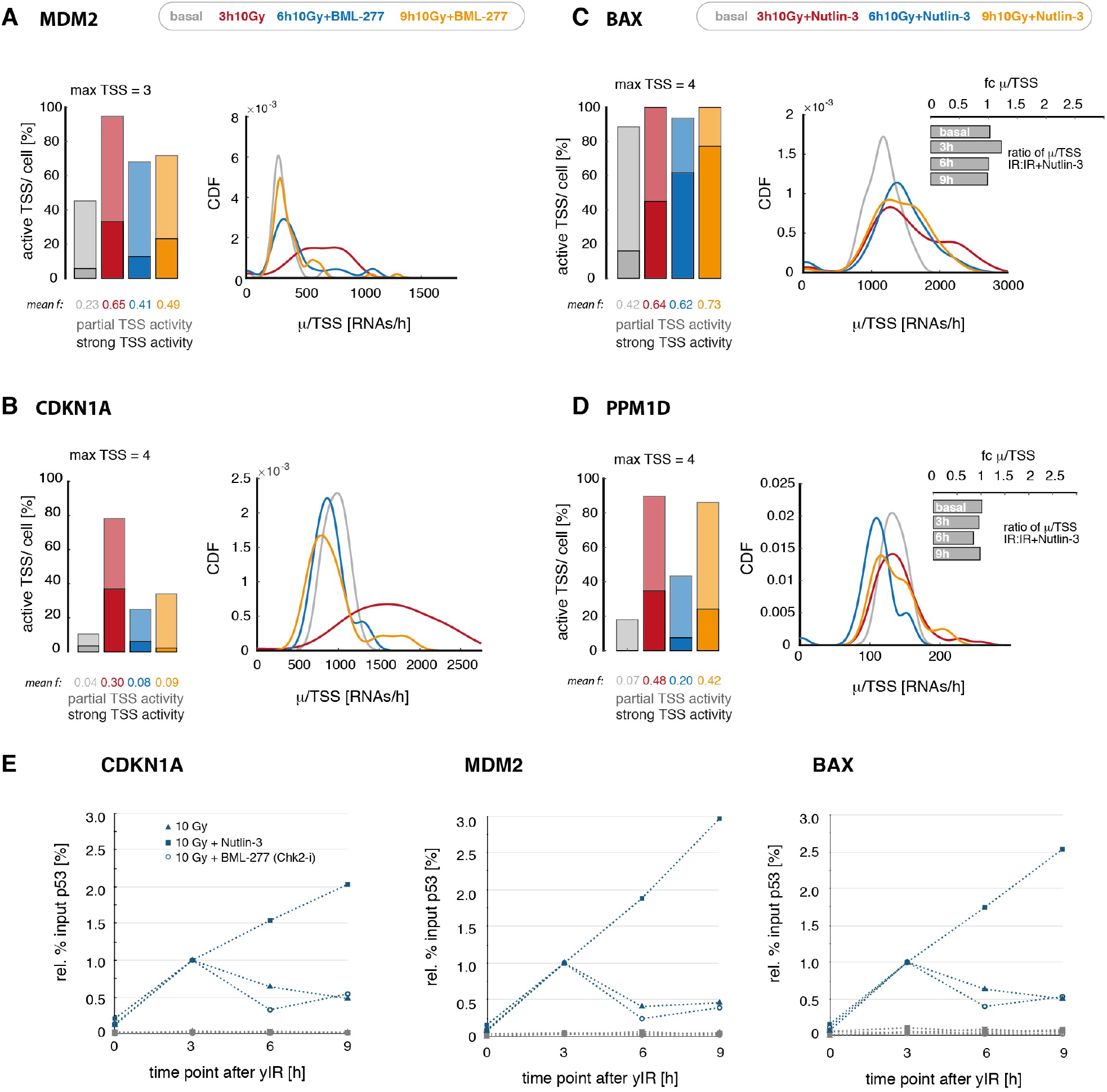
Quantification of bursting kinetics of target genes upon modulated p53 dynamics. **A/B** We quantified the fraction of active promoters (left panel) and burst size per TSS (right panel) of MDM2 (A) and CDKN1A (B) after BML-277 treatment inducing transient p53 dynamics. As both target genes were grouped in the transient archetype, no obvious changes in promoter activity were observed. Shaded colors: partially active cells, solid colors: cells in which a majority of TSS are actively transcribing, average fractions of active promoters are indicated at the bottom of each panel (mean f). **C/D** We measured the fraction of active promoters (left panel) and burst size per TSS (right panel) upon sequential Nutlin-3 treatment inducing sustained p53 dynamics for BAX (C) and PPM1D (D). Relative burst size per TSS compared to previous experiments with pulsatile p53 dynamics are indicated as well (inset, fold change calculated for each time point after IR (fc μ/TSS)). For BAX, a target gene that we grouped into the sustained archetype, both transcription parameters remain high. Interestingly in the contrary to our observations to target genes that change their gene expression mode in response to Nutlin-3 treatment, the burst size of BAX transcription does not change strongly. Surprisingly, the same hold true for PPM1D, a gene that we grouped in to the pulsatile archetype. Shaded colors: partially active cells, solid colors: cells in which a majority of TSS are actively transcribing, average fractions of active promoters are indicated at the bottom of each panel (mean f). **E** To measure relative p53 binding at different target gene promoters in perturbed and unperturbed cells, we performed ChIP experiments in the context of Nutlin-3 and BML-277 treatment for all p53 target genes at the indicated time points after 10Gy IR - CDKN1A (left), MDM2 (center) and BAX (right) (we could not detect any p53 binding at the PPM1D promoter in repeated experiments). Results were normalized to the 3h time point post IR. Grey symbols indicate the corresponding IgG control. Interestingly, pulsatile p53 and inhibition of the second p53 pulse both lead to a gradual decrease of p53 binding after the first peak, Nutlin-3 treatment increases p53 binding at these promoters and may thereby contribute to the observed increase in promoter activity.

**Figure EV4.**
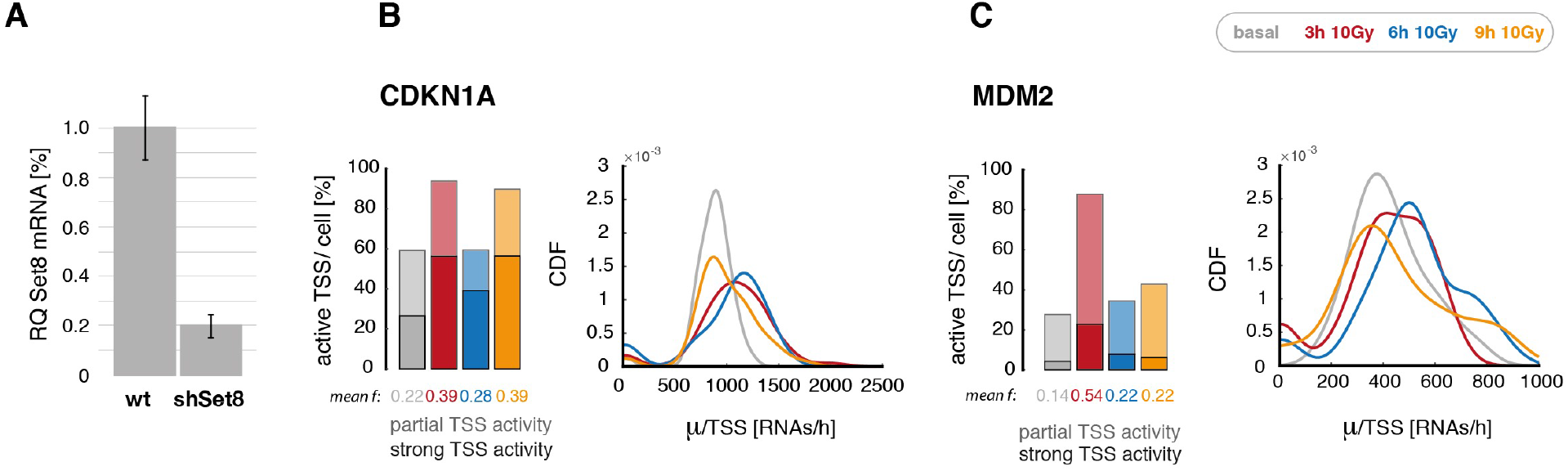
Bursting kinetics of target genes upon Set8 knock down. **A** The p53-K382 methylase Set8 was down-regulated in a clonal stable A549 cell line expressing a corresponding shRNA. Transcript levels were measured in wild type and knock-down cells by qRT-PCR. Mean levels and standard deviation from technical triplicates are indicated. **B/C** Promoter activity of CDKN1A (B) and MDM2 (C) were quantified in Set8 knock-down cells after 10 Gray yIR. We observed that the burst size per TSS (left panel, [RNAs/h]) in Set8 knock down cells is similar to wild type cells, while the fraction of active promoters is increased (right panel, [%]). Shaded colors: partially active cells, solid colors: cells in which a majority of TSS are actively transcribing, average fractions of active promoters are indicated at the bottom of each panel (mean f).

